# Prediction of Threonine-Tyrosine Kinase Receptor-Ligand Unbinding Kinetics with Multiscale Milestoning and Metadynamics

**DOI:** 10.1101/2024.08.01.606239

**Authors:** Lane W. Votapka, Anupam Anand Ojha, Naoya Asada, Rommie E. Amaro

## Abstract

Accurately describing protein-ligand binding and unbinding kinetics remains challenging. Computational calculations are difficult and costly, while experimental measurements often lack molecular detail and can be unobtainable. Here we extend our multiscale milestoning method, Simulation-Enabled Estimation of Kinetics Rates (SEEKR), with metadynamics molecular dynamics simulations to yield accurate small molecule drug residence times. Using the pharmaceutically relevant threonine-tyrosine kinase (TTK) and eight long-residence-time (tens of seconds to hours) inhibitors, we demonstrate accurate prediction of absolute and rank-ordered ligand residence times and free energies of binding.

**TOC Graphic:** 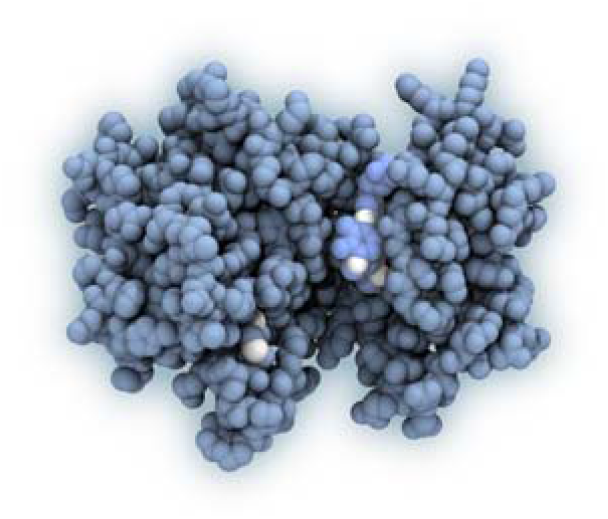

## Introduction

### Background

Biological systems involve a vast array of complex intermolecular interactions. The binding and unbinding of a ligand-receptor pair can be described by 1) thermodynamics, which considers equilibrium quantities like the free energy of binding (ΔG_bind_) and the equilibrium constant (K_D_), and 2) kinetics, which considers time-dependent quantities like the rate constant of binding (k_on_), the rate constant of unbinding (k_off_), and the residence time (1/k_off_).^1^ Knowledge of these quantities is desirable for predicting a drug’s efficacy^2–5^ and optimizing specificity in drug discovery^6–8^. While experimental measurements of binding/unbinding kinetics and thermodynamics are possible, accurate and efficient computational predictions remain attractive. SEEKR is a multiscale, simulation-based, enhanced sampling method used to characterize the kinetics and thermodynamics of a variety of binding/unbinding systems^9–15^.

Metadynamics (metaD) is an enhanced sampling method that estimates the free energy landscapes of complicated systems^16–20^. It accelerates the exploration of the free energy landscape by introducing a CV-based history-dependent bias potential that evolves over time, searching for new metastable states during MD simulations^17,21^. Previously, SEEKR used steered molecular dynamics (SMD) for generating starting structures; in this work, we use the GPU-accelerated well-tempered metaD implementation within SEEKR. Threonine-tyrosine kinase (TTK), also known as monopolar spindle 1 (MPS1) is frequently overexpressed in highly proliferative cancers, making it a promising drug target for human breast cancer^22^. A recent study revealed that residence time, rather than potency (IC_50_), has a stronger correlation with anti-proliferative activity^23^. Improving the kinetic properties of TTK inhibitors is essential, but the binding and unbinding mechanism is still unclear.

**Figure 1:**
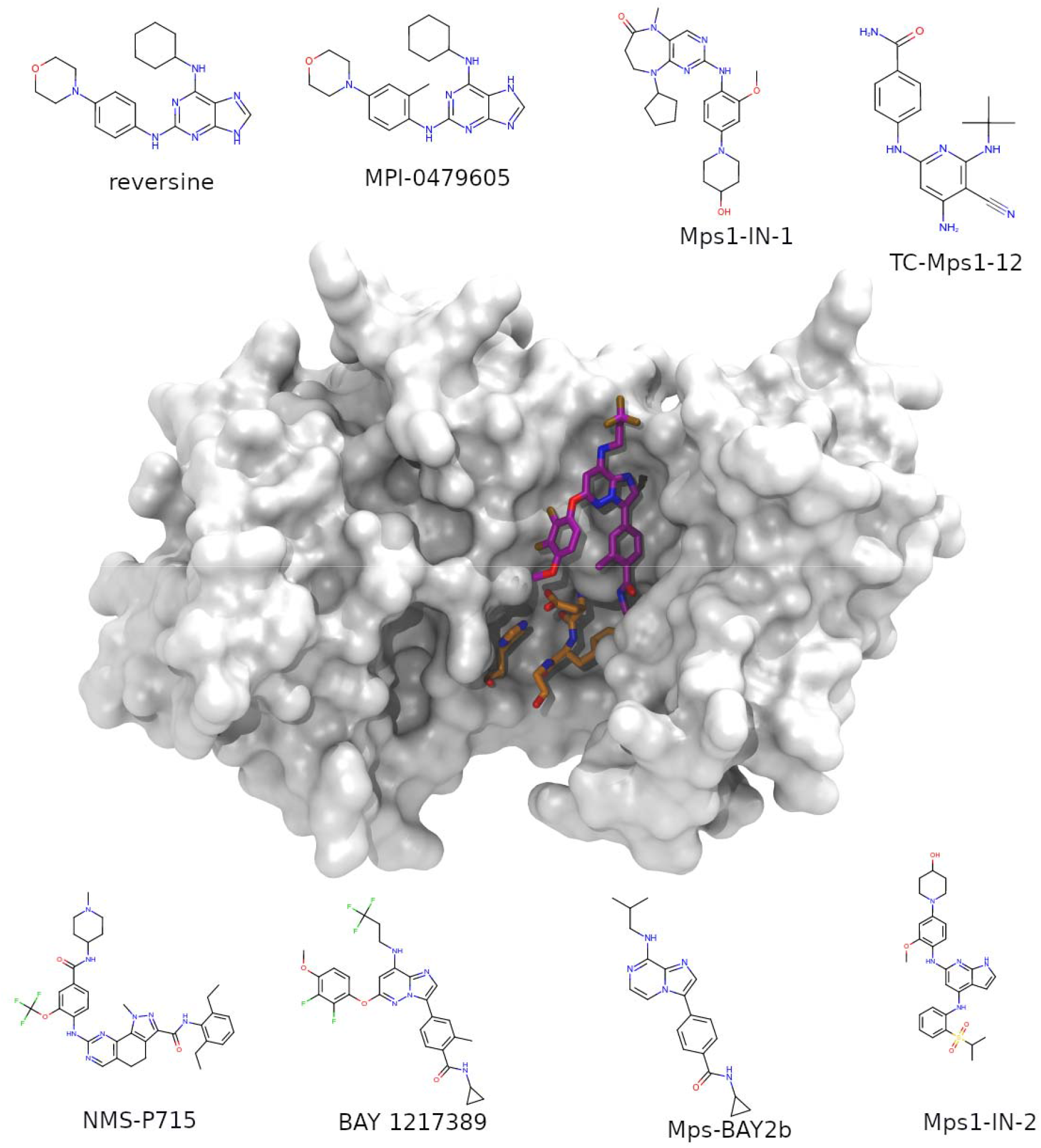
TTK structure, prominent features, and ligands. In the center, the TTK protein is shown as a translucent white surface, taken from the X-ray crystal structure with PDB ID: 5NAD^23^. It is bound to compound BAY 1217389, shown with purple carbons. Within the protein, the Mg^2+^ binding (DFG) loop, composed of ASP664, PHE665, and GLY666 is colored with orange carbons, as well as HIS645, which makes a significant interaction with ASP 664.

We ran SEEKR on eight TTK systems with experimental residence times available. Details of the computational methods, benchmarks, and simulation costs are in the SI. SEEKR requires generated starting structures for simulations within the milestoning framework. Previously, this was done with steered molecular dynamics (SMD), which uses a harmonic restraint moving at a constant velocity along the CV to sample starting structures. In this study, we explore the use of metaD, which gradually fills the energy landscape with Gaussian boosts, allowing the ligand to gradually sample steep energy landscape portions. This comparison of initial sampling approaches within SEEKR - using SMD versus metaD - can be found in Figure 2 (Exact numerical values can be found in Tables S1 and S2).

**Figure 2:**
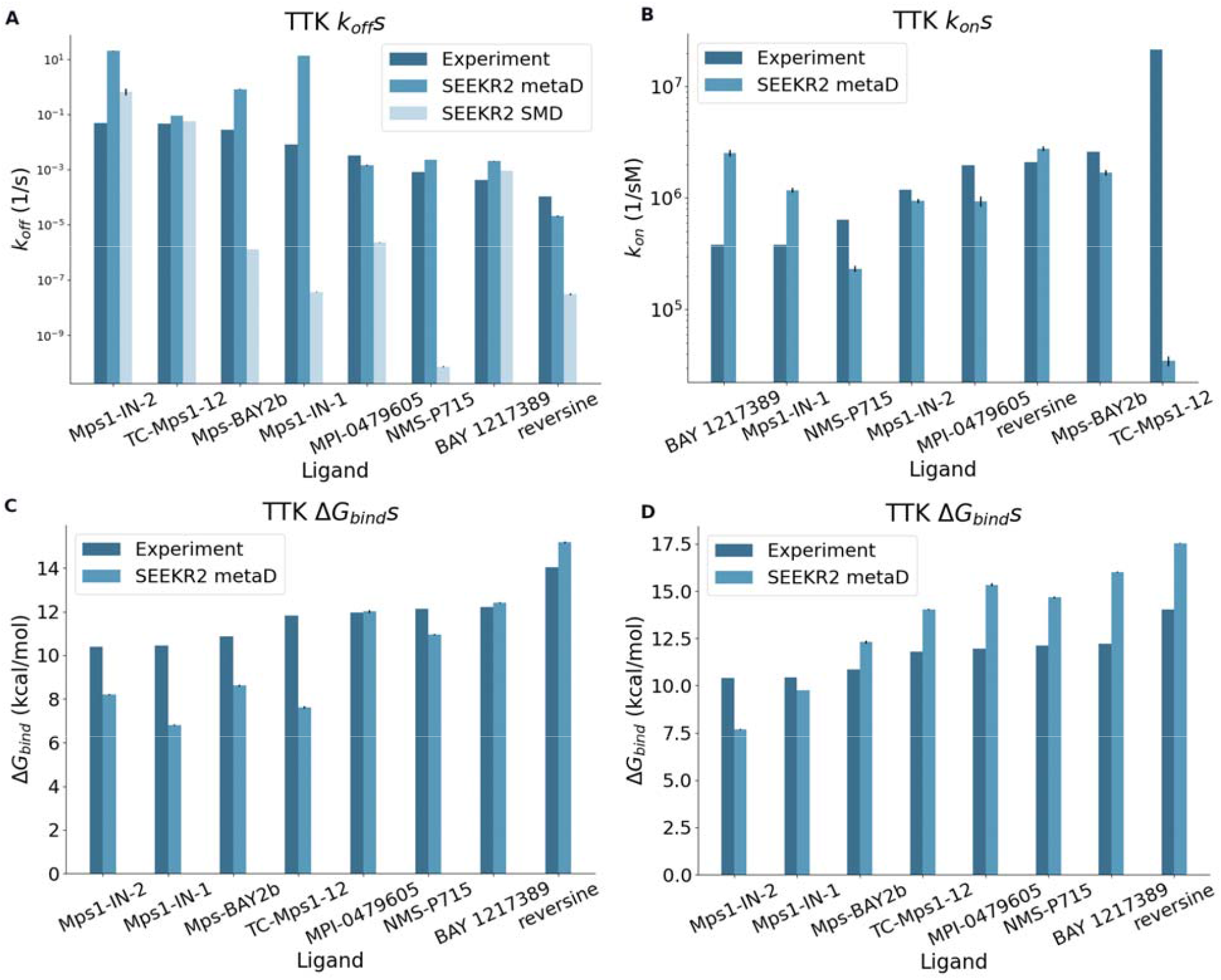
TTK k_off_ rate constants (A), k_on_ rate constants (B), and ΔG_bind_ (C) for known inhibitors. A) Results for k_off_ show that SEEKR2 with metaD produced the best results (medium blue). In contrast, SMD (light blue) produces sub-optimal results. B) Predictions within a single order of magnitude of experiment were found for all TTK system k_on_ predictions except TC-Mps1-12. The reason for the one outlier is uncertain, and additional analysis and investigations would need to be performed to identify the reason for it. C) These were computed by using the equation ΔG_bind_ = k_B_T·ln(k_off_/k_on_). Clearly, the biggest deviation in TC-Mps1-12 was caused by the highly incorrect k_on_ obtained for that system. D) These were computed by the difference in free energies of the MMVT anchors between the bound state and the pre-defined ‘escaped’ state at the mouth of the binding site (as defined in the supplementary information). While there is a large systematic bias caused by the missing solvation energy of a true separation between ligand and receptor, the trend is very good - generating almost perfect ranking.

**Figure 3:**
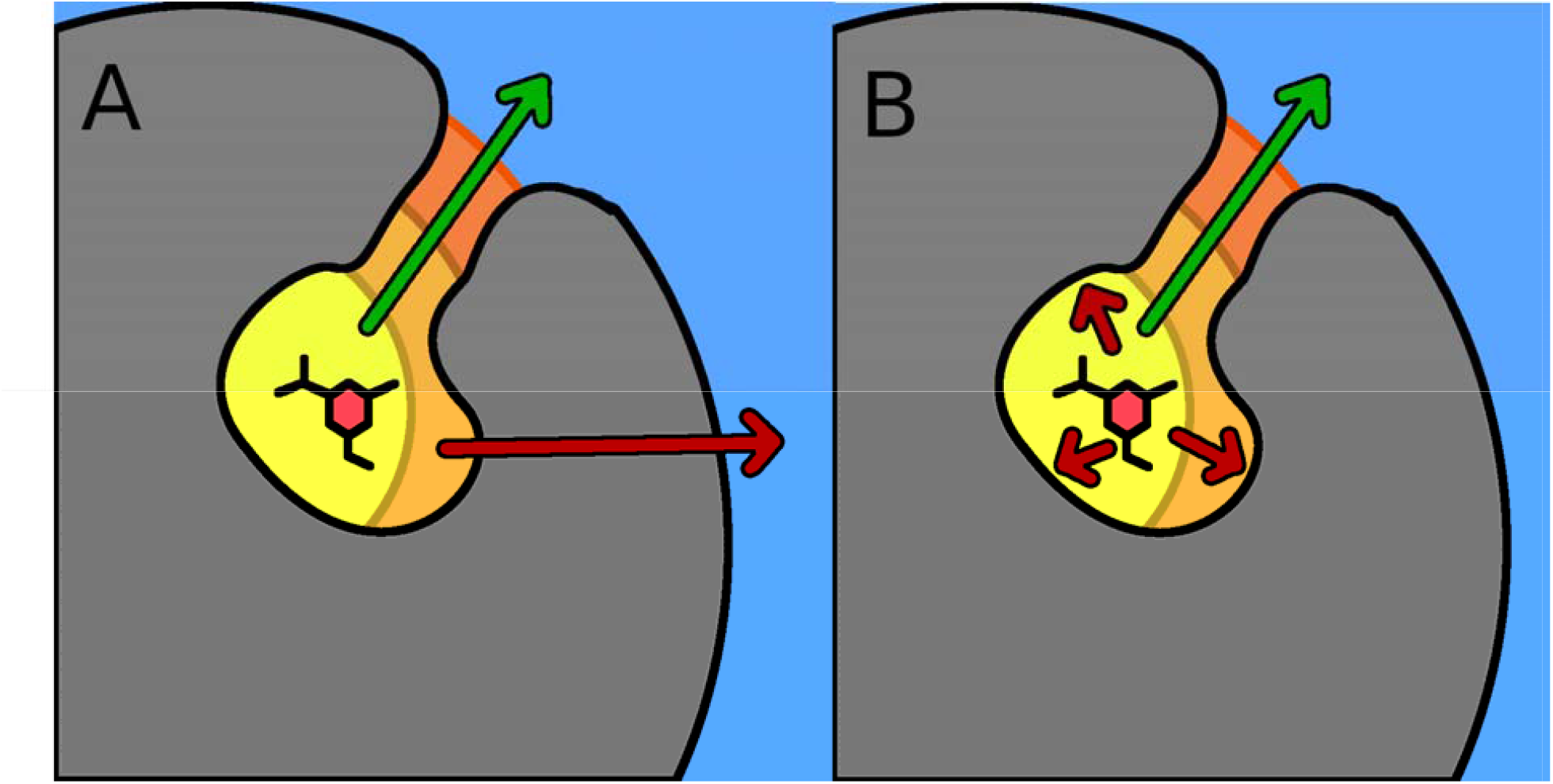
Cartoon schematic comparing exploration of correct and incorrect unbinding pathways using SMD (panel A) and metaD (panel B). In both panels, the green arrow represents the correct unbinding path, while the incorrect unbinding paths are red arrow(s). In panel A, SMD might explore the correct unbinding path, but if the ligand gets pushed along the incorrect unbinding path, up against an interior surface of the binding site, the inability of SMD to “back up” causes the ligand to get pushed through the interior surface. In contrast, in panel B, metaD can also sample the correct unbinding path, but if the incorrect unbinding path is explored, the surface of the binding site will stop the ligand, and the metaD sampling algorithm will allow the ligand to backtrack.

SEEKR obtains a Kendall’s tau of 0.64 and a mean absolute log_10_ error of 1.2 for ranking of absolute unbinding kinetics using metaD, and a Kendall’s tau of 0.50 with a mean absolute log_10_ error of 3.1 for the same with SMD and QMrebind. Values for k_on_ and ΔG_bind_ were also computed for the TTK system and are reported in Figure 2B, 2C, and 2D. The Kendall’s tau for this series of k_on_ values was found to be -0.18, and the mean absolute log_10_ of the error was computed to be 0.66; for this series of ΔG_bind_ values (computed from k_B_T·ln(k_off_/k_on_)), the Kendall’s tau was 0.71, with a mean absolute error of 1.9 kcal/mol. When the ΔG_bind_ values were computed from the relative free energies of MMVT anchors, it gives a Kendall’s tau of 0.93, and a mean absolute error of 2.5 kcal/mol. (Exact numerical values for k_off_ values can be found in Table S1 k_on_, and ΔG_bind_ values can be found in Table S2).

In addition to the method of kinetics prediction, the choice and preparation of system structure files for the receptor and ligand are crucial^24–27^, including the selection of high-quality experimentally-determined structures, ensuring physical realism (matching experimental conditions like temperature, pressure, ion concentrations, and pH), carefully considering protonation states of ligands and receptor residues, and performing adequate minimization and equilibration to ensure that the system is modeled near a thermodynamic equilibrium state. The choice of force field parameters for SEEKR calculations is also critical. We use the QMrebind method^14^ to redefine the partial charges of the ligand in the bound state, which works well for the TTK system (Figure 2), particularly affecting the partial charges on a tert-butyl side group (Figure S1). Other force field parameters, such as ligand torsions, may also benefit from refinement - a topic we do not address here. All the partial charges, both before and after using QMrebind to reparametrize them, can be found in Table S3.

Challenges arise when modeling halogens^28^ due to the “sigma hole”, producing a positive partial charge opposite the covalent bond with chlorine, bromine, or iodine atoms. In particular, inaccuracies have been observed when the halogen interacts with Lewis bases. However, fluorine’s small sigma hole can be adequately described with a point charge^29^, as shown by the successful application of SEEKR to the TTK 5NAD and 2X9E systems, which contain many fluorines.

After system preparation, choosing the CV, and providing force field parameters, the SEEKR MMVT algorithm populates each Voronoi cell with starting structures to run simulations and gather milestoning times and statistics. The HIDR tool within SeekrTools (https://github.com/seekrcentral/seekrtools.git) facilitates this by exploring the entire reaction coordinate span and saving potential starting structures. We found that metaD provides superior starting structures compared to other methods. Previously, SMD was commonly used for generating starting structures for SEEKR. However, more complex ligands with large, flexible structures lead to artificial tearing and unfolding of the protein. SMD sometimes pushed the ligand along the wrong unbinding pathway or through the binding site’s interior surface, resulting in inaccurate SEEKR kinetics predictions (Figure 9A).

While adverse effects of SMD can sometimes be mitigated by carefully reselecting the atoms defining the binding site, alternative sampling algorithms like MetaD allow the ligand to backtrack and explore multiple exit pathways. MetaD works well for this purpose, unlike SMD, which constrains the ligand to a particular CV value and can push it through the binding site’s interior surface if stuck.

However, using metaD to generate starting structures has its challenges. If the Gaussian height is too large, it can disrupt the molecular system, affecting the accuracy of the SEEKR calculation (data not shown). Conversely, if the Gaussian heights are too small, generating starting structures can be time-consuming and resource-intensive. The choice of CV is also nontrivial and can complicate practical use, especially for less-studied receptors^30–32^.

Convergence is important for SEEKR calculations to determine if additional sampling is necessary or if an incorrect physical description impacts accuracy. Convergence plots in Figure S2 show that metaD starting structures converge better than SMD, likely because metaD samples structures with lower energy, closer to the true unbinding pathway.

In this work, we expand SEEKR’s capabilities and demonstrate its success in estimating binding and unbinding kinetics for the TTK system and eight small molecules with residence times from seconds to hours. The combined use of metaD for initial structure generation and quantum-mechanically reparametrized ligands significantly enhances SEEKR calculations.

SEEKR2 open-source software is available at GitHub: https://github.com/seekrcentral/seekr2.git, and SeekrTools, which runs metaD and SMD, at https://github.com/seekrcentral/seekrtools.git. Data for the systems in this study can be obtained from https://doi.org/10.6075/J0571C7G.

## Acknowledgements

The authors thank Steffen Wolf, University of Freiburg, for his insightful and valuable discussions. The authors express their gratitude to Mac Kevin E. Braza for his assistance with the figures. Thanks to Jeff Wagner for helpful discussions about force field parameterization as well as assistance with testing, continuous integration, and GitHub. A. A. O. acknowledges the support of the Merkin Fellowship from the University of California San Diego. All simulations were performed using the Popeye computing cluster at the San Diego Supercomputing Center (SDSC) and the Delta supercomputing system at the National Center for Supercomputing Applications (NCSA), University of Illinois Urbana-Champaign. The authors acknowledge support from NSF Advanced Cyberinfrastructure Coordination Ecosystem: Services and Support (ACCESS) CHE060063. The authors thank the Laboratory for Medicinal Chemistry Research, Shionogi and CO. Ltd, Osaka, Japan, for their collaborative efforts and contributions.

