## Supplementary Information for "Prediction of Threonine-Tyrosine Kinase Receptor-Ligand Unbinding Kinetics with Multiscale Milestoning and Metadynamics"

**Supplementary Information for Enhanced Receptor-Ligand Unbinding Kinetics with Metadynamics, Quantum Corrections and Multiscale Milestoning**

All SEEKR-derived results within cells of Table S1 and S2 are highlighted by color according to their closeness to experimental values. For $k_{on}$ and $k_{off}$, green indicates a SEEKR result within a single order of magnitude difference from experiment, yellow indices within two orders of magnitude, orange within three orders of magnitude, and red is a SEEKR result that deviates more than three orders of magnitude from the experimental value. For $\Delta G_{bind}$ , green indicates a SEEKR result within one kcal/mol of difference from experiment, yellow indices within two kcal/mol, orange within three kcal/mol, and red is a SEEKR result that deviates more than three kcal/mol from the experimental value

### Computational Methods

All SEEKR calculations were performed using the crystal structures 5LJJ[1], 5N84[2], 5N93[2], 5N7V[2] 5NAD[2], 2X9E[3], 3GFW[4], and 3H9F[4]. For the preparation, protein preparation wizard in Schrödinger 2021 suites[5] was used to add hydrogen atoms and consider the protonation state of the system. Protonation states of TTK were initially found with the Schrodinger Epik program within Maestro, and later confirmed and adjusted using PDB2PQR[6]. TTK experimental kinetics and thermodynamics values were obtained from a study that recognized the importance of residence time to the cellular potency for the TTK target[2]. The protein and solvent portions of the system were parameterized using the LEAP program within the AmberTools 20[7] software suite. We created the topologies and initial coordinate files using the Amber ff19SB force field[8] for the protein with the GAFF force field[9] and AM1BCC charge were assigned for the ligand, though then reassigned with QMrebind[10] following energy minimization. TIP4P-Ew was used for water parameters[11]. Within QMrebind, the protein-ligand complexes were partitioned into three regions: the high-level QM region, comprised of the ligand itself; the low-level QM2 region, comprised of complete residues with any atoms residing within 5 Å of any of the ligand atoms, and finally, the MM region, which contained all the rest of the atoms. The QM region was modeled using the cc-pVTZ basis set with an MP2 method for the inner QM region. The QM2 region was modeled with the GFN2-XTB semiempirical tight-binding method[12]. Structures were then minimized and equilibrated using OpenMM[13]. CPPtraj was used to auto-image the solvent around the complex[14].

All systems contained a site center-of-mass to ligand center-of-mass (COM-COM) distance CV. Unless specified otherwise, all active sites atom definitions (to characterize the COM-COM distance CV) were defined by examining the native bound structure, selecting all alpha carbon atoms within 3 Å of the COM of the ligand, and then gradually removing them one at a time, or adding more alpha carbons within 6 Å of the COM of the ligand, until the set of alpha carbons defining the site had a COM within 0.5 Å of the ligand COM. Following this, milestone spacings were chosen in a system-dependent manner, typically choosing closer spacings for regions of the CV where the ligand is within the site and wider spacings for CV regions where the ligand is out of the site or immersed in the solvent. Details on CV definitions and milestoning placements for each system are provided in the supplementary information tables S4 and S5. Following parameterization and equilibration, SMD simulations were run within HIDR of SeekrTools for 1μs for all systems with a restraint force constant of $9 \cdot 10^4$

kJ/(mol·nm$^2$) , and starting structures were saved every time the anchor point was crossed. metaD simulations were run within HIDR with a Gaussian width of 0.05 nm, a Gaussian height of 1.0 kJ/mol, and a bias factor of 10. Binding site definitions for the TTK system can be found in Table S4.

Milestone placements for system TTK can be found in Table S5. The system structures were saved upon the last time that the simulations crossed each anchor point along the relevant CV. Although the metaD Gaussian height parameter was system-dependent, all metaD simulations used a sigma value of 0.5 Å and a bias factor of 10. Following this, SEEKR2 runs were performed in parallel for 500 ns per anchor using MMVT milestoning, which uses OpenMM as its MD engine[13], and all systems were subsequently analyzed using the native SEEKR2 and SeekrTools analysis programs. PQR files were prepared directly from AMBER parameter/topology files output from LEAP, using the MM partial charges and the radii from the mbondi2 set for the atomic charges and radii, respectively. The Browndye programs were used to prepare and run all BD simulations within the SEEKR framework[15]. Biomolecular images were generated with VMD[16].

The mean absolute error (MAE) for any set of data was computed with: $\sum_i \left| x_{i,calc} - x_{i,exp} \right|$ where $x_{i,calc}$ and $x_{i,exp}$ are the SEEKR-calculated and experimental quantities, respectively, that are being compared. In some cases, differences between calculated and experimental quantities exceeded many orders of magnitude, so the logarithm base ten of the calculated and experimental data, $log_{10}(x_{i,calc})$ and $log_{10}(x_{i,exp})$ were used to obtain the mean absolute $\log_{10}$ error. The $\Delta G_{bind}$ values computed from the relative free energies of MMVT anchors were found using the equation $k_B T \cdot \ln(\pi_{escaped}/\pi_{bound})$, where $\pi_\alpha$ is the stationary probability of anchor α, and is found using MMVT theory[17].

**Benchmarking and Simulation Costs**

We chose to simulate each anchor for 500 ns, and since each system possessed 23 simulated anchors, each different system required 11,500 ns - a total of 184,000 ns required for the MMVT simulations of all sixteen systems. The SMD simulations were run for 1000 ns per system, requiring a total of 8000 ns MD for the eight SMD systems. (The metaD simulations were relatively short - in the tens of ns - so we neglect these simulation costs). The anchors were benchmarked to run at approximately 315 ns/day on a V100 GPU, and since all anchors for a system could be run synchronously on a large GPU cluster, the entire SEEKR2 calculation for a system (using metaD) could be completed in approximately 40 hours, using ~875 GPU hours per metaD system. To run SMD required an additional 1000 ns per system, which needed approximately 76 additional hours, for a final cost of ~950 GPU hours for a SMD SEEKR2 calculation. The eight metaD systems and eight SMD systems therefore required approximately 15,330 GPU hours to complete.

#### Table S1 - Unbinding kinetics for TTK systems

| TTK System (PDB ID) | Experimental $k_{off}$ ($s^{-1}$) | SEEKR $k_{off}$ ($s^{-1}$) using 1000 ns SMD, with QMrebind | SEEKR $k_{off}$ ($s^{-1}$) using MetaD, with QMrebind (correct protonation) |
|---|---|---|---|
| Reversine (5LJJ) | $1.00 \cdot 10^{-4}$ | $(3.1 \pm 0.2) \cdot 10^{8}$ | $(2.03 \pm 0.09) \cdot 10^{-5}$ |
| MPI-0479605 (5N7V) | $3.40 \cdot 10^{-3}$ | $(2.26 \pm 0.08) \cdot 10^{-6}$ | $(1.46 \pm 0.08) \cdot 10^{-3}$ |
| TC-Mps1-12 (5N93) | $4.70 \cdot 10^{-2}$ | $(5.70 \pm 0.05) \cdot 10^{-2}$ | $(8.99 \pm 0.06) \cdot 10^{-2}$ |
| BAY 1217389 (5NAD) | $4.10 \cdot 10^{-4}$ | $(9.2 \pm 0.1) \cdot 10^{-4}$ | $(2.01 \pm 0.03) \cdot 10^{-3}$ |
| Mps-BAY2b (5N84) | $2.80 \cdot 10^{-2}$ | $(1.30 \pm 0.02) \cdot 10^{-6}$ | $(8.3 \pm 0.2) \cdot 10^{-1}$ |
| NMS-P715 (2X9E) | $8.33 \cdot 10^{-4}$ | $(7.1 \pm 0.3) \cdot 10^{-11}$ | $(2.29 \pm 0.02) \cdot 10^{-3}$ |
| Mps1-IN-1 (3GFW) | $8.33 \cdot 10^{-3}$ | $(3.7 \pm 0.1) \cdot 10^{-8}$ | $(1.36 \pm 0.02) \cdot 10^{1}$ |
| Mps1-IN-2 (3H9F) | $5.0 \cdot 10^{-2}$ | $(6.6 \pm 1.8) \cdot 10^{-1}$ | $(2.00 \pm 0.09) \cdot 10^{1}$ |

#### Table S2 - Binding kinetics and free energies for TTK systems

| Compound | Experimental $k_{on}$ (1/s·M) | SEEKR $k_{on}$ (1/s·M) | Experimental $\Delta G_{bind}$ (kcal/mol) | SEEKR $\Delta G_{bind}$ from $K_D$ (kcal/mol) | SEEKR $\Delta G_{bind}$ from milestoning (kcal/mol) |
|---|---|---|---|---|---|
| Reversine (5LJJ) | $2.08 \cdot 10^{6}$ | $(2.8 \pm 0.1) \cdot 10^{6}$ | 14.0 | $15.18 \pm 0.04$ | $17.54 \pm 0.03$ |
| MPI-0479605 (5N7V) | $1.96 \cdot 10^{6}$ | $(9. \pm 1.) \cdot 10^{5}$ | 12.0 | $12.00 \pm 0.07$ | $15.34 \pm 0.07$ |
| TC-Mps1-12 (5N93) | $2.16 \cdot 10^{7}$ | $(3.4 \pm 0.4) \cdot 10^{4}$ | 11.8 | $7.61 \pm 0.06$ | $14.033 \pm 0.003$ |
| BAY 1217389 (5NAD) | $3.79 \cdot 10^{5}$ | $(2.5 \pm 0.2) \cdot 10^{6}$ | 12.2 | $12.41 \pm 0.04$ | $16.01 \pm 0.02$ |
| Mps-BAY2b (5N84) | $2.60 \cdot 10^{6}$ | $(1.69 \pm 0.09) \cdot 10^{6}$ | 10.9 | $8.60 \pm 0.04$ | $12.31 \pm 0.08$ |
| NMS-P715 (2X9E) | $6.41 \cdot 10^{5}$ | $(2.3 \pm 0.2) \cdot 10^{5}$ | 12.1 | $10.95 \pm 0.04$ | $14.67 \pm 0.05$ |
| Mps1-IN-1 (3GFW) | $3.79 \cdot 10^{5}$ | $(1.17 \pm 0.06) \cdot 10^{6}$ | 10.4 | $6.79 \pm 0.03$ | $9.764 \pm 0.006$ |
| Mps1-IN-2 (3H9F) | $1.19 \cdot 10^{6}$ | $(9.4 \pm 0.4) \cdot 10^{5}$ | 10.4 | $8.20 \pm 0.04$ | $7.68 \pm 0.03$ |

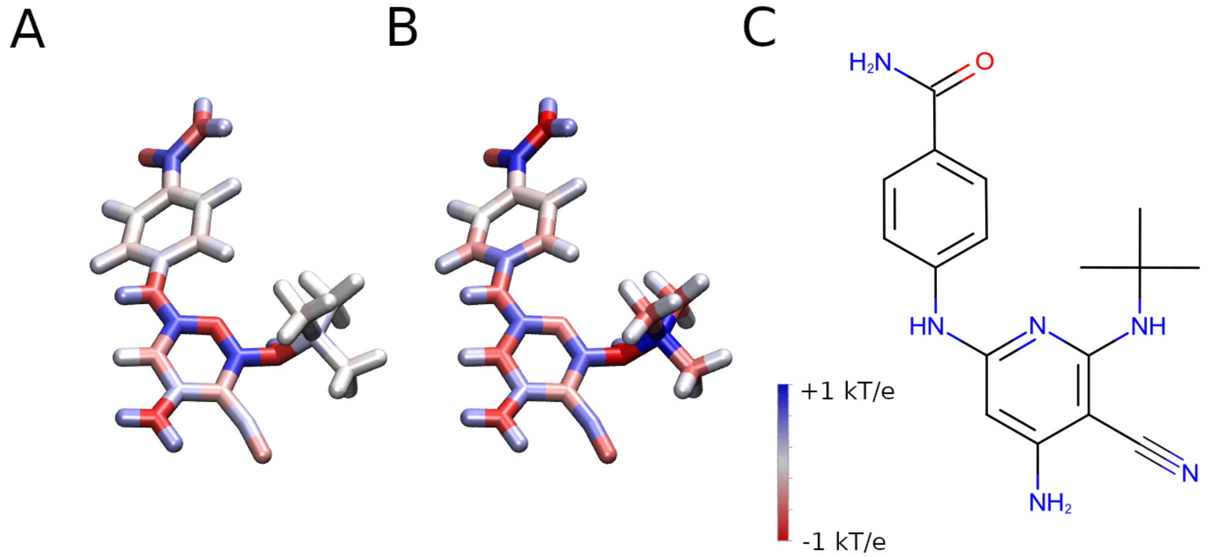

**Figure S1:** Reparametrization of charges using QMrebind on compound TC-Mps1-12. (A) Charges of the compound assigned by ordinary AM1BCC in GAFF. (B) Charges on the same compound reparametrized with QMrebind - note the changes to the charges on the tert-butyl group. (C) 2D representation of the compound for reference.

### Table S3 - TTK system ligand charge reparametrization from AM1BCC charges to QMrebind charges

All charges in units of proton charge

| Reversine (5LJJ) charges | | | | MPI-0479605 (5N7V) charges | | | | TC-Mps1-12 (5N93) charges | | |
|---|---|---|---|---|---|---|---|---|---|---|
| Atom Name | AM1BCC charge | QMrebind charge | | Atom Name | AM1BCC charge | QMrebind charge | | Atom Name | AM1BCC charge | QMrebind charge |
| C1 | -0.0944 | -0.5019 | | C6 | -0.0939 | -0.1837 | | C12 | -0.1061 | -0.6005 |
| H1 | 0.049 | 0.129 | | H2 | 0.0542 | 0.0599 | | H7 | 0.0459 | 0.1131 |
| H2 | 0.049 | 0.1117 | | H3 | 0.0542 | 0.072 | | H8 | 0.0459 | 0.1471 |
| C2 | -0.0749 | 0.1215 | | C7 | -0.0814 | -0.1837 | | H9 | 0.0459 | 0.1355 |
| H3 | 0.0429 | 0.028 | | H4 | 0.041 | 0.0742 | | C11 | 0.2452 | 1.0591 |
| H4 | 0.0429 | -0.011 | | H5 | 0.041 | 0.0449 | | C13 | -0.1061 | -0.6519 |
| C3 | -0.0804 | -0.1794 | | C8 | -0.0794 | 0.0622 | | H10 | 0.0459 | 0.129 |
| H5 | 0.0407 | 0.0728 | | H6 | 0.0407 | 0.0063 | | H11 | 0.0459 | 0.1412 |
| H6 | 0.0407 | 0.0686 | | H7 | 0.0407 | 0.0047 | | H12 | 0.0459 | 0.1592 |
| C4 | -0.0749 | -0.0702 | | C9 | -0.0814 | -0.0745 | | C14 | -0.1061 | -0.669 |
| H7 | 0.0429 | 0.0434 | | H8 | 0.041 | 0.0158 | | H13 | 0.0459 | 0.1677 |
| H8 | 0.0429 | 0.0572 | | H9 | 0.041 | 0.0289 | | H14 | 0.0459 | 0.1638 |
| C5 | -0.0944 | -0.1845 | | C10 | -0.0939 | -0.1424 | | H15 | 0.0459 | 0.1442 |
| H9 | 0.049 | 0.0245 | | H10 | 0.0542 | 0.0387 | | N3 | -0.8519 | -1.0205 |
| H10 | 0.049 | 0.0595 | | H11 | 0.0542 | 0.0336 | | H6 | 0.4338 | 0.396 |
| C6 | 0.2185 | 0.6276 | | C11 | 0.2115 | 0.5768 | | C7 | 0.7822 | 0.6644 |
| H11 | 0.0837 | -0.0157 | | H12 | 0.0917 | -0.0061 | | N4 | -0.813 | -0.5276 |
| N1 | -0.8409 | -0.6746 | | N1 | -0.8249 | -0.8405 | | C6 | -0.3393 | -0.4166 |
| H12 | 0.4428 | 0.3065 | | H13 | 0.4238 | 0.3329 | | C8 | 0.2558 | 0.5165 |
| C7 | 0.7802 | 0.358 | | C3 | 0.7712 | 0.6129 | | N1 | -0.4088 | -0.6085 |
| C8 | -0.2602 | 0.1889 | | N2 | -0.823 | -0.6151 | | C9 | 0.3206 | 0.4544 |
| N3 | -0.5341 | -0.5931 | | C2 | -0.2402 | -0.0113 | | N2 | -0.8592 | -0.8209 |
| C9 | 0.3907 | 0.1114 | | N3 | -0.5341 | -0.5477 | | H3 | 0.4318 | 0.4188 |
| N4 | -0.387 | -0.4939 | | C4 | 0.3897 | 0.1152 | | H4 | 0.4318 | 0.3752 |
| C10 | 0.4878 | 0.5836 | | N4 | -0.389 | -0.4469 | | C10 | -0.4453 | -0.7095 |
| N5 | -0.79 | -0.9598 | | H14 | 0.3157 | 0.4094 | | H5 | 0.143 | 0.273 |
| H14 | 0.3157 | 0.4028 | | H1 | 0.0708 | 0.185 | | C5 | 0.7442 | 0.6614 |
| H13 | 0.0708 | 0.1757 | | C1 | 0.4838 | 0.628 | | N5 | -0.7303 | -0.6564 |
| N2 | -0.816 | -0.6249 | | N5 | -0.79 | -0.9883 | | H16 | 0.4298 | 0.3847 |
| C11 | 0.9159 | 0.8978 | | C5 | 0.9159 | 0.7459 | | C3 | 0.1716 | 0.5202 |
| N6 | -0.6983 | -0.7445 | | N6 | -0.6973 | -0.3601 | | C2 | -0.1615 | -0.4007 |
| H15 | 0.4328 | 0.439 | | H15 | 0.4318 | 0.3318 | | H1 | 0.1475 | 0.2142 |
| C12 | 0.1026 | 0.449 | | C12 | 0.0976 | -0.1646 | | C15 | -0.073 | -0.0297 |
| C15 | -0.111 | -0.248 | | C16 | -0.073 | -0.1707 | | H17 | 0.146 | 0.1685 |

| | | | | | | | | | | |
|---|---|---|---|---|---|---|---|---|---|---|
| C16 | -0.157 | -0.4389 | | C17 | -0.175 | -0.4483 | | C1 | -0.1636 | -0.1788 |
| H19 | 0.135 | 0.2334 | | H21 | 0.132 | 0.2414 | | C17 | 0.6737 | 0.8746 |
| H18 | 0.1425 | 0.2023 | | H20 | 0.151 | 0.1887 | | O1 | -0.6131 | -0.656 |
| C13 | -0.111 | -0.3334 | | C13 | -0.0833 | 0.6771 | | N6 | -0.675 | -1.1166 |
| H16 | 0.1425 | 0.2102 | | C14 | -0.0608 | -0.5909 | | H19 | 0.3115 | 0.4573 |
| C14 | -0.157 | -0.1987 | | H16 | 0.047 | 0.1237 | | H20 | 0.3115 | 0.4752 |
| H17 | 0.135 | 0.1539 | | H17 | 0.047 | 0.1947 | | C16 | -0.073 | -0.1384 |
| C17 | 0.1236 | 0.2423 | | H18 | 0.047 | 0.0968 | | H18 | 0.146 | 0.1933 |
| N7 | -0.628 | -0.3323 | | C15 | -0.145 | -0.7716 | | C4 | -0.1615 | -0.3634 |
| C18 | 0.1478 | 0.2669 | | H19 | 0.137 | 0.2236 | | H2 | 0.1475 | 0.1573 |
| H20 | 0.0474 | -0.0087 | | C18 | 0.1266 | 0.5981 | | | | |
| H21 | 0.0474 | -0.0447 | | N7 | -0.622 | -0.3619 | | | | |
| C19 | 0.1239 | -0.0323 | | C19 | 0.1478 | 0.0271 | | | | |
| H22 | 0.0559 | 0.0937 | | H22 | 0.0597 | 0.0546 | | | | |
| H23 | 0.0559 | 0.0562 | | H23 | 0.0597 | 0.0259 | | | | |
| O1 | -0.4146 | -0.4248 | | C20 | 0.1029 | 0.2857 | | | | |
| C20 | 0.1239 | -0.0378 | | H24 | 0.0519 | 0.0031 | | | | |
| H24 | 0.0559 | 0.1137 | | H25 | 0.0519 | 0.0016 | | | | |
| H25 | 0.0559 | 0.0806 | | O1 | -0.4136 | -0.5335 | | | | |
| C21 | 0.1478 | 0.1507 | | C21 | 0.1029 | 0.2136 | | | | |
| H26 | 0.0474 | 0.0612 | | H26 | 0.0519 | 0.0344 | | | | |
| H27 | 0.0474 | 0.0319 | | H27 | 0.0519 | 0.0572 | | | | |
| | | | | C22 | 0.1478 | -0.1933 | | | | |
| | | | | H28 | 0.0597 | 0.1054 | | | | |
| | | | | H29 | 0.0597 | 0.103 | | | | |
| BAY 1217389 (5NAD) | | | | Mps-BAY2b (5N84) | | | | NMS-P715 (2X9E) | | |
| Atom Name | AM1BCC charge | QMrebind charge | | Atom Name | AM1BCC charge | QMrebind charge | | Atom Name | AM1BCC charge | QMrebind charge |
| F2 | -0.1059 | -0.1409 | | C19 | -0.0926 | -0.4492 | | C22 | 0.115 | -0.194 |
| C15 | 0.1409 | 0.0041 | | H18 | 0.0369 | 0.0772 | | C23 | -0.131 | -0.143 |
| C9 | 0.0259 | 0.3167 | | H19 | 0.0369 | 0.0911 | | H23 | 0.090 | 0.121 |
| F1 | -0.1259 | -0.1757 | | H20 | 0.0369 | 0.113 | | H24 | 0.090 | 0.145 |
| C10 | 0.1251 | -0.0224 | | C18 | -0.1017 | 0.5717 | | H21 | 0.105 | 0.132 |
| O2 | -0.3209 | -0.2711 | | C20 | -0.0926 | -0.7372 | | H22 | 0.105 | 0.138 |
| C11 | 0.1107 | -0.0816 | | H21 | 0.0369 | 0.1475 | | N8 | -0.686 | 0.062 |
| H7 | 0.054 | 0.0645 | | H22 | 0.0369 | 0.1404 | | C24 | 0.084 | -0.528 |
| H8 | 0.054 | 0.0453 | | H23 | 0.0369 | 0.1717 | | H25 | 0.106 | 0.211 |
| H9 | 0.054 | 0.1462 | | H17 | 0.0637 | -0.048 | | H26 | 0.106 | 0.245 |

| | | | | | | | | | | |
|---|---|---|---|---|---|---|---|---|---|---|
| C12 | -0.146 | -0.0819 | | C17 | 0.1988 | -0.1145 | | H27 | 0.106 | 0.191 |
| H10 | 0.165 | 0.1799 | | H15 | 0.0667 | 0.0821 | | H20 | 0.447 | 0.355 |
| C13 | -0.081 | -0.4551 | | H16 | 0.0667 | 0.1163 | | C21 | 0.115 | -0.297 |
| H11 | 0.162 | 0.2402 | | N5 | -0.8219 | -0.4903 | | H18 | 0.105 | 0.170 |
| C14 | 0.0401 | 0.4388 | | H14 | 0.4388 | 0.3555 | | H19 | 0.105 | 0.175 |
| O1 | -0.3414 | -0.4088 | | C14 | 0.7132 | 0.5858 | | C20 | -0.131 | -0.071 |
| C1 | 0.6406 | 0.7315 | | C15 | -0.1936 | 0.0988 | | H16 | 0.090 | 0.132 |
| N1 | -0.5728 | -0.688 | | N4 | -0.2706 | -0.6372 | | H17 | 0.090 | 0.094 |
| C3 | -0.3703 | -0.5121 | | C16 | 0.0236 | 0.1169 | | C19 | 0.105 | 0.282 |
| H1 | 0.171 | 0.2491 | | H13 | 0.186 | 0.1634 | | H15 | 0.108 | 0.076 |
| C2 | 0.2936 | -0.0211 | | N3 | -0.742 | -0.6065 | | N7 | -0.586 | -0.842 |
| N4 | -0.7486 | -0.2909 | | C13 | 0.4082 | 0.2372 | | H14 | 0.319 | 0.395 |
| C6 | 0.2088 | -0.0277 | | H12 | 0.0371 | 0.142 | | C18 | 0.696 | 0.886 |
| C7 | -0.1374 | -0.2293 | | C12 | -0.2586 | -0.789 | | O3 | -0.593 | -0.696 |
| C8 | 0.6209 | 0.8003 | | H11 | 0.177 | 0.3257 | | C15 | -0.200 | -0.157 |
| F3 | -0.2386 | -0.3084 | | N2 | 0.0737 | 0.7065 | | C16 | -0.061 | -0.235 |
| F4 | -0.2386 | -0.2709 | | C11 | -0.1283 | -0.5371 | | C17 | -0.171 | -0.190 |
| F5 | -0.2386 | -0.2616 | | C8 | 0.0102 | 0.565 | | H13 | 0.169 | 0.199 |
| H5 | 0.0962 | 0.116 | | C9 | -0.132 | -0.4579 | | H12 | 0.131 | 0.153 |
| H6 | 0.0962 | 0.0792 | | C10 | -0.0875 | 0.0508 | | C14 | -0.023 | -0.297 |
| H3 | 0.0827 | 0.1201 | | H10 | 0.1485 | 0.1112 | | H11 | 0.179 | 0.178 |
| H4 | 0.0827 | 0.1074 | | H9 | 0.143 | 0.1795 | | C13 | -0.001 | 0.385 |
| H12 | 0.4378 | 0.3338 | | C7 | -0.132 | -0.4507 | | O2 | -0.367 | -0.408 |
| C16 | -0.1273 | 0.4219 | | H8 | 0.143 | 0.2363 | | C25 | 0.813 | 0.883 |
| N3 | -0.2706 | -0.5404 | | C6 | -0.0875 | -0.0389 | | F1 | -0.209 | -0.218 |
| C5 | 0.0046 | -0.1139 | | H7 | 0.1485 | 0.1637 | | F2 | -0.209 | -0.201 |
| H2 | 0.188 | 0.2245 | | C5 | -0.1416 | -0.2261 | | F3 | -0.209 | -0.284 |
| N2 | 0.2302 | 0.4146 | | C1 | 0.6777 | 0.7165 | | C12 | 0.251 | 0.198 |
| C4 | -0.0693 | -0.3496 | | O1 | -0.5981 | -0.6415 | | N6 | -0.727 | -0.628 |
| C17 | 0.0042 | 0.6003 | | N1 | -0.5419 | -0.6215 | | H10 | 0.480 | 0.423 |
| C23 | -0.109 | -0.5479 | | H1 | 0.3115 | 0.3135 | | C11 | 0.872 | 0.962 |
| C21 | -0.0253 | 0.3668 | | C2 | 0.0537 | 0.2501 | | N5 | -0.747 | -0.702 |
| C22 | -0.0718 | -0.6059 | | H2 | 0.1227 | 0.0823 | | C8 | 0.554 | 0.556 |
| H15 | 0.058 | 0.1979 | | C3 | -0.1539 | -0.1529 | | N4 | -0.710 | -0.822 |
| H16 | 0.058 | 0.1978 | | H3 | 0.0739 | 0.1093 | | C10 | 0.461 | 0.384 |
| H17 | 0.058 | 0.1645 | | H4 | 0.0739 | 0.1436 | | H9 | 0.044 | 0.127 |
| H18 | 0.152 | 0.2505 | | C4 | -0.1539 | -0.5603 | | C9 | -0.308 | -0.346 |
| C18 | -0.127 | -0.4662 | | H5 | 0.0739 | 0.2 | | C2 | -0.025 | -0.152 |
| H13 | 0.14 | 0.1788 | | H6 | 0.0739 | 0.1942 | | H3 | 0.065 | 0.098 |

| | | | | | | | | | | |
|---|---|---|---|---|---|---|---|---|---|---|
| C19 | -0.105 | -0.0977 | | | | | | H4 | 0.065 | 0.099 |
| H14 | 0.14 | 0.2002 | | | | | | C1 | -0.016 | -0.002 |
| C20 | -0.1396 | -0.3167 | | | | | | H1 | 0.085 | 0.042 |
| C24 | 0.6817 | 0.9213 | | | | | | H2 | 0.085 | 0.081 |
| O3 | -0.5971 | -0.7766 | | | | | | C3 | -0.134 | 0.045 |
| N5 | -0.5439 | -0.623 | | | | | | C4 | -0.220 | -0.228 |
| H19 | 0.3135 | 0.3949 | | | | | | N1 | 0.147 | 0.347 |
| C25 | 0.0537 | 0.0554 | | | | | | C7 | 0.040 | -0.295 |
| H20 | 0.1217 | 0.1797 | | | | | | H6 | 0.067 | 0.105 |
| C26 | -0.1544 | -0.4717 | | | | | | H7 | 0.067 | 0.158 |
| H21 | 0.0737 | 0.1776 | | | | | | H8 | 0.067 | 0.111 |
| H22 | 0.0737 | 0.1806 | | | | | | N2 | -0.486 | -0.478 |
| C27 | -0.1544 | -0.1481 | | | | | | C5 | 0.252 | -0.023 |
| H23 | 0.0737 | 0.0965 | | | | | | C6 | 0.601 | 0.797 |
| H24 | 0.0737 | 0.1083 | | | | | | O1 | -0.588 | -0.697 |
| | | | | | | | | N3 | -0.462 | -0.415 |
| | | | | | | | | H5 | 0.324 | 0.322 |
| | | | | | | | | C26 | 0.024 | -0.136 |
| | | | | | | | | C27 | -0.064 | 0.282 |
| | | | | | | | | C34 | -0.043 | -0.149 |
| | | | | | | | | C35 | -0.093 | -0.155 |
| | | | | | | | | H38 | 0.037 | 0.052 |
| | | | | | | | | H39 | 0.037 | 0.024 |
| | | | | | | | | H40 | 0.037 | 0.018 |
| | | | | | | | | H36 | 0.054 | 0.099 |
| | | | | | | | | H37 | 0.054 | 0.083 |
| | | | | | | | | C28 | -0.128 | -0.403 |
| | | | | | | | | H28 | 0.139 | 0.189 |
| | | | | | | | | C29 | -0.123 | 0.094 |
| | | | | | | | | H29 | 0.135 | 0.151 |
| | | | | | | | | C30 | -0.128 | -0.687 |
| | | | | | | | | H30 | 0.139 | 0.306 |
| | | | | | | | | C31 | -0.064 | 0.189 |
| | | | | | | | | C32 | -0.043 | 0.137 |
| | | | | | | | | H31 | 0.054 | 0.001 |
| | | | | | | | | H32 | 0.054 | 0.063 |
| | | | | | | | | C33 | -0.093 | -0.334 |
| | | | | | | | | H33 | 0.037 | 0.076 |
| | | | | | | | | H34 | 0.037 | 0.110 |

| | | | | | | | | | | |
|---|---|---|---|---|---|---|---|---|---|---|
| | | | | | | | | H35 | 0.037 | 0.076 |
| Mps1-IN-1 (3GFW) | | | | Mps1-IN-2 (3H9F) | | | | | | |
| Atom Name | AM1BCC charge | QMrebind charge | | Atom Name | AM1BCC charge | QMrebind charge | | | | |
| C1 | 0.197 | -0.208 | | C4 | 0.198 | 0.084 | | | | |
| H1 | 0.044 | 0.161 | | H2 | 0.046 | 0.094 | | | | |
| H2 | 0.044 | 0.030 | | H3 | 0.046 | 0.065 | | | | |
| C2 | -0.125 | 0.131 | | C5 | -0.109 | -0.413 | | | | |
| H3 | 0.062 | -0.037 | | H4 | 0.062 | 0.128 | | | | |
| H4 | 0.062 | 0.000 | | H5 | 0.062 | 0.139 | | | | |
| C3 | 0.142 | 0.291 | | C6 | 0.144 | 0.490 | | | | |
| O1 | -0.596 | -0.676 | | O1 | -0.601 | -0.763 | | | | |
| H6 | 0.399 | 0.369 | | H7 | 0.400 | 0.376 | | | | |
| H5 | 0.078 | -0.054 | | H6 | 0.036 | -0.026 | | | | |
| C4 | -0.125 | -0.063 | | C7 | -0.109 | -0.051 | | | | |
| H7 | 0.062 | 0.005 | | H8 | 0.062 | -0.038 | | | | |
| H8 | 0.062 | 0.022 | | H9 | 0.062 | 0.060 | | | | |
| C5 | 0.197 | 0.043 | | C8 | 0.198 | 0.034 | | | | |
| H9 | 0.044 | 0.000 | | H10 | 0.046 | -0.007 | | | | |
| H10 | 0.044 | 0.080 | | H11 | 0.046 | 0.088 | | | | |
| N1 | -0.654 | -0.032 | | N3 | -0.651 | -0.272 | | | | |
| C6 | 0.191 | 0.080 | | C9 | 0.185 | 0.203 | | | | |
| C10 | -0.220 | -0.411 | | C13 | -0.217 | -0.399 | | | | |
| C11 | -0.059 | -0.202 | | C14 | -0.045 | -0.108 | | | | |
| H16 | 0.150 | 0.200 | | H17 | 0.154 | 0.185 | | | | |
| H15 | 0.138 | 0.258 | | H16 | 0.138 | 0.206 | | | | |
| C7 | -0.246 | -0.273 | | C10 | -0.240 | -0.355 | | | | |
| H11 | 0.143 | 0.116 | | H12 | 0.142 | 0.143 | | | | |
| C8 | 0.134 | 0.289 | | C11 | 0.128 | 0.285 | | | | |
| O2 | -0.333 | -0.342 | | O2 | -0.341 | -0.370 | | | | |
| C9 | 0.116 | -0.026 | | C12 | 0.118 | 0.059 | | | | |
| H12 | 0.047 | 0.011 | | H13 | 0.046 | 0.078 | | | | |
| H13 | 0.047 | 0.097 | | H14 | 0.046 | 0.099 | | | | |
| H14 | 0.047 | 0.109 | | H15 | 0.046 | -0.004 | | | | |
| C12 | 0.032 | 0.153 | | C15 | 0.050 | 0.055 | | | | |
| N2 | -0.684 | -0.558 | | N4 | -0.668 | -0.536 | | | | |
| H17 | 0.425 | 0.368 | | H18 | 0.439 | 0.362 | | | | |
| C13 | 0.669 | 0.841 | | C16 | 0.866 | 0.847 | | | | |

| | | | | | | |
|---|---|---|---|---|---|---|
| N3 | -0.743 | -1.023 | | N2 | -0.766 | -0.551 |
| C15 | 0.481 | 0.820 | | N1 | -0.737 | -0.877 |
| N4 | -0.294 | -0.561 | | C3 | 0.450 | 0.434 |
| C16 | -0.116 | -0.114 | | H1 | 0.034 | 0.130 |
| C17 | -0.163 | -0.417 | | C2 | -0.235 | -0.289 |
| C18 | -0.268 | -0.256 | | C1 | 0.662 | 0.458 |
| H21 | 0.166 | 0.210 | | N6 | -0.738 | -0.373 |
| H20 | 0.169 | 0.209 | | C21 | 0.206 | 0.358 |
| H19 | 0.309 | 0.450 | | C22 | -0.093 | -0.123 |
| C14 | -0.340 | -0.821 | | C23 | -0.080 | -0.043 |
| H18 | 0.156 | 0.262 | | C24 | -0.080 | -0.061 |
| C19 | 0.245 | 0.558 | | C25 | -0.093 | -0.069 |
| N5 | -0.653 | -0.641 | | H33 | 0.051 | 0.039 |
| H22 | 0.480 | 0.425 | | H34 | 0.051 | 0.012 |
| C20 | 0.315 | 0.351 | | H31 | 0.042 | 0.012 |
| C21 | -0.218 | -0.244 | | H32 | 0.042 | 0.056 |
| H23 | 0.158 | 0.155 | | H29 | 0.042 | -0.002 |
| C22 | -0.032 | -0.122 | | H30 | 0.042 | 0.069 |
| H24 | 0.135 | 0.176 | | H27 | 0.051 | 0.019 |
| C23 | -0.199 | -0.238 | | H28 | 0.051 | -0.003 |
| H25 | 0.142 | 0.164 | | H26 | 0.073 | -0.003 |
| C24 | -0.006 | -0.090 | | C20 | 0.193 | 0.017 |
| H26 | 0.157 | 0.175 | | H24 | 0.052 | 0.030 |
| C25 | -0.460 | -0.157 | | H25 | 0.052 | 0.153 |
| S1 | 1.366 | 0.969 | | C19 | -0.159 | -0.392 |
| O3 | -0.663 | -0.566 | | H22 | 0.080 | 0.165 |
| O4 | -0.663 | -0.632 | | H23 | 0.080 | 0.106 |
| C26 | -0.261 | 0.239 | | C18 | 0.714 | 0.822 |
| C28 | -0.071 | -0.633 | | O3 | -0.636 | -0.764 |
| H31 | 0.049 | 0.206 | | N5 | -0.363 | -0.206 |
| H32 | 0.049 | 0.147 | | C17 | 0.082 | -0.342 |
| H33 | 0.049 | 0.177 | | H19 | 0.049 | 0.160 |
| H27 | 0.113 | 0.132 | | H20 | 0.049 | 0.175 |
| C27 | -0.071 | -0.510 | | H21 | 0.049 | 0.148 |
| H28 | 0.049 | 0.150 | | | | |
| H29 | 0.049 | 0.106 | | | | |
| H30 | 0.049 | 0.176 | | | | |

### Figure S2 - Convergence of $k_{off}$ for TTK Systems

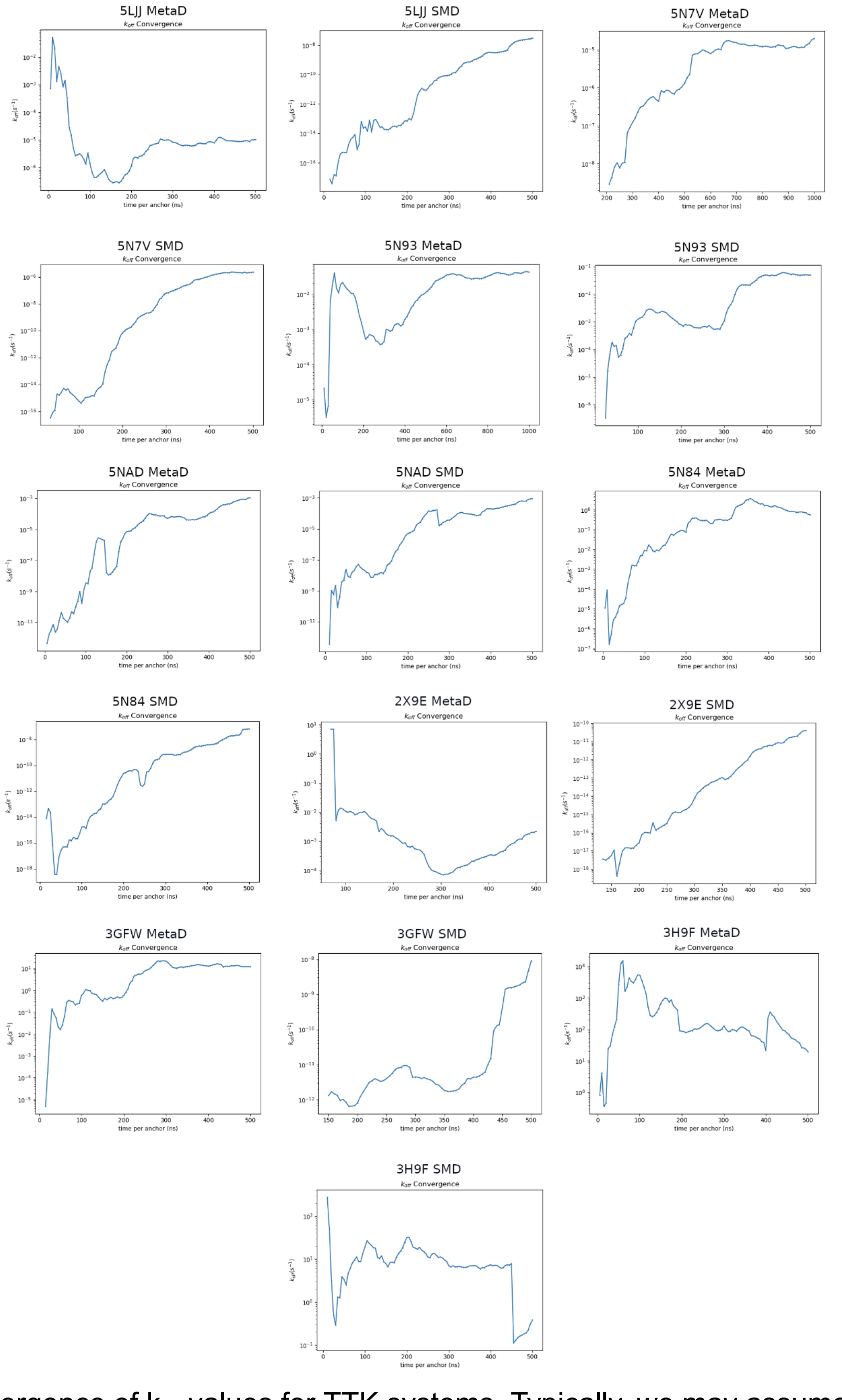

Figure S2: Convergence of $k_{off}$ values for TTK systems. Typically, we may assume that a system is converged if the $k_{off}$ fluctuates within only an order of magnitude during the last half of the simulation. Using this criteria, many SMD simulations are not fully converged within the 1000ns of the SEEKR simulations, with the possible exception of 5NAD SMD. While MetaD simulations were more likely to converge (probably due to more energetically-favorable chosen starting structures), not all were converged by the end of the simulation - 5N84 for instance.

**Table S4 - TTK system binding site definitions:**

Sites were chosen by selecting alpha carbons (C$\alpha$) of all residues with any atoms within 0.3 nm of any atoms of the ligand, then individually adding nearby residue C$\alpha$s or removing residue C$\alpha$s from the selection until the center of mass (COM) of the site fell within 0.05 nm of the ligand COM.

| Ligand (PDBID) | Site Atoms (Atom Name - Residue Name - Residue Index) |
|---|---|
| Reversine (5LJJ) | CA-ILE-531, CA-VAL-539, CA-ALA-551, CA-TYR-604, CA-GLY-605, CA-ILE-607, CA-ASP-608, CA-ALA-651, CA-LEU-654, CA-ILE-663, CA-THR-675 |
| MPI-0479605 (5N7V) | CA-VAL-539, CA-GLN-541, CA-ALA-551, CA-CYS-604, CA-ILE-607, CA-ASP-608, CA-LEU-654, CA-ILE-663, CA-MET-671 |
| TC-Mps1-12 (5N93) | CA-VAL-539, CA-GLN-541, CA-ALA-551, CA-CYS-604, CA-ILE-607, CA-ASP-608, CA-LEU-654, CA-ILE-663, CA-MET-671 |
| BAY 1217389 (5NAD) | CA-LYS-529, CA-ILE-531, CA-GLN-541, CA-ALA-551, CA-LYS-553, CA-ILE-586, CA-MET-602, CA-CYS-604, CA-GLY-605, CA-ASP-608, CA-LYS-649, CA-ALA-651, CA-LEU-654, CA-ILE-663, CA-MET-671, CA-PRO-673 |
| Mps-BAY2b (5N84) | CA-ILE-531, CA-VAL-539, CA-LYS-553, CA-LEU-575, CA-CYS-604, CA-ILE-607, CA-ASP-608, CA-ILE-663, CA-VAL-684 |
| NMS-P715 (2X9E) | CA-LEU-528, CA-LYS-529, CA-GLN-530, CA-ILE-531, CA-GLY-532, CA-SER-533, CA-GLY-534, CA-SER-537, CA-LYS-538, CA-VAL-539, CA-GLN-541, CA-ALA-551, CA-LYS-553, CA-MET-602, CA-GLU-603, CA-CYS-604, CA-GLY-605, CA-ASN-606, CA-ILE-607, CA-ASP-608, CA-SER-611, CA-TRP-612, CA-LYS-614, CA-LYS-615, CA-ALA-651, CA-ASN-652, CA-LEU-654, CA-ILE-663 |
| Mps1-IN-1 (3GFW) | CA-LYS-529, CA-ILE-531, CA-GLY-532, CA-SER-533, CA-SER-537, CA-VAL-539, CA-GLN-541, CA-ALA-551, CA-LYS-553, CA-ILE-586, CA-GLU-603, CA-CYS-604, CA-GLY-605, CA-ASN-606, CA-ILE-607, CA-ASP-608, CA-SER-611, CA-LYS-614, CA-LYS-615, CA-ALA-651, CA-ASN-652, CA-LEU-654, CA-PRO-673, CA-THR-676 |
| Mps1-IN-2 (3H9F) | CA-ILE-531, CA-LYS-553, CA-MET-602, CA-CYS-604, CA-GLY-605, CA-ILE-607, CA-SER-611, CA-LEU-654, CA-PRO-673, CA-TPO-675 |

**Table S5 - TTK system anchor radii**

For the BAY 1217389 and Mps-BAY2b systems, because of the large size of those ligands, a set of anchors that ended at a further distance from the site was observed to better represent the unbound state. The radii highlighted in **bold** show the state that was considered “escaped” for $k_{off}$ calculations (chosen by visual inspection of SEEKR trajectories - the distance when the ligand could traverse to regions of the protein surface other than the immediate opening of the binding site). The higher anchors were still used for $k_{on}$ calculations.

| **Ligand (PDBID)** | **Anchor radii (nm)** |
|---|---|
| Reversine (5LJJ) | 0.1500, 0.2250, 0.3000, 0.3750, 0.4500, 0.5250, 0.6000, 0.6750, 0.7750, 0.8750, 1.0000, 1.1000, 1.2000, 1.3000, 1.4000, 1.5000, 1.6000, 1.7000, 1.8000, **1.9000,** 2.0000, 2.2000, 2.4000, 2.6000 |
| MPI-0479605 (5N7V) | 0.1500, 0.2250, 0.3000, 0.3750, 0.4500, 0.5250, 0.6000, 0.7000, 0.8000, 0.9000, 1.0000, 1.1000, 1.2000, 1.3000, 1.4000, 1.5000, 1.6000, 1.7000, 1.8000, **1.9000,** 2.0000, 2.2000, 2.4000, 2.6000 |
| TC-Mps1-12 (5N93) | 0.1500, 0.2250, 0.3000, 0.3750, 0.4500, 0.5250, 0.6000, 0.7000, 0.8000, 0.9000, 1.0000, 1.1000, 1.2000, 1.3000, 1.4000, 1.5000, **1.6000**, 1.7000, 1.8000, 1.9000**,** 2.0000, 2.2000, 2.4000, 2.6000 |
| BAY 1217389 (5NAD) | 0.1500, 0.2250, 0.3000, 0.3750, 0.4500, 0.5250, 0.6000, 0.7000, 0.8000, 0.9000, 1.0000, 1.1000, 1.2000, 1.3000, 1.4000, 1.5000, 1.6000, 1.7000, 1.8000, 2.0000, 2.2000, 2.4000, **2.6000**, 2.8000 |
| Mps-BAY2b (5N84) | 0.1500, 0.2250, 0.3000, 0.3750, 0.4500, 0.5250, 0.6000, 0.7000, 0.8000, 0.9000, 1.0000, 1.1000, 1.2000, 1.3000, 1.4000, 1.5000, 1.6000, 1.7000, **1.8000**, 2.0000, 2.2000, 2.4000, 2.6000, 2.8000 |
| NMS-P715 (2X9E) | 0.1500, 0.2250, 0.3000, 0.3750, 0.4500, 0.5250, 0.6000, 0.7000, 0.8000, 0.9000, 1.0000, 1.1000, 1.2000, 1.3000, 1.4000, 1.5000, 1.6000, 1.7000, 1.8000, 2.0000, 2.2000, 2.4000, **2.6000**, 2.8000 |
| Mps1-IN-1 (3GFW) | 0.1500, 0.2250, 0.3000, 0.3750, 0.4500, 0.5250, 0.6000, 0.7000, 0.8000, 0.9000, 1.0000, 1.1000, 1.2000, 1.3000, 1.4000, 1.5000, 1.6000, 1.7000, 1.8000, 2.0000, 2.2000, 2.4000, **2.6000**, 2.8000 |
| Mps1-IN-2 (3H9F) | 0.1500, 0.2250, 0.3000, 0.3750, 0.4500, 0.5250, 0.6000, 0.7000, 0.8000, 0.9000, 1.0000, 1.1000, 1.2000, 1.3000, 1.4000, 1.5000, 1.6000, 1.7000, 1.8000, 2.0000, 2.2000, 2.4000, **2.6000**, 2.8000 |
